# iMOUSE - Reforming the Strategy of Refinement and Reduction for indispensable laboratory animal-based studies in translational research

**DOI:** 10.1101/2023.08.06.552188

**Authors:** Maciej Łaz, Mirko Lampe, Isaac Connor, Dmytro Shestachuk, Marcel Ludwig, Ursula Müller, Oliver F. Strauch, Nadine Suendermann, Stefan Lüth, Janine Kah

## Abstract

Considering the intricate nature of biological processes within organisms, it is undeniable that relying solely on in vitro-generated primary-cell-like cultures or organ-like products in preclinical and basic research is insufficient to replace animal-based studies fully. This limitation is particularly significant when considering the regulations enforced by legislative assemblies worldwide. The necessity of animal-based studies to approve chemicals and medications. In contradiction, European countries aim to banish animal-based studies. Therefore, we must understand the impact of the data refinement and experiment replacement strategy we will introduce here.

This project **aimed** to revolutionize data acquisition in animal-based studies by transforming manual observation into a reliable digital process. Reliable digital data will be generated by having the potential to reduce human bias by simply reducing human interaction. Additionally, reducing human interaction will reduce the severity levels due to stress reduction, fulfilling the 3R principles.

**Therefore, the first goal was** to develop and implement a scalable, stable, running, and remotely accessible camera-based monitor system (the iMouse solution). At the same time, the target was to develop a retrofit solution (DigiFrame) for existing home-cage systems, not interfering with the regular workflow in animal facilities. **As a result**, we developed a digital monitoring system, named iMouseTV platform based on existing open-source software, allowing users to observe, record, share, and review animal-based studies within the home cage anytime from anywhere, reducing the stress level for the animals. Our system’s first Proof of concept ran for over two years at the LIV in Hamburg. We also investigated an effective way to reduce data generation by setting up specific zones for detecting the motion of choice (e.g., drinking, food intake). The data sets can be stored, shared, and reviewed by users and refined by algorithms aiming to recognize the dedicated motions of the animals automatically. The implementation of the ML algorithms allows the iMouse solution to recognize whether an individual mouse was drinking and for how long and store results in the annotated video file and graph format. However, the identification and continuous tracking of the species is still in progress.

In conclusion, we established a scalable human-independent monitoring and recording system, which can be implemented into the existing structures of institutions and companies without changing handling processes, to monitor animals and observe them by getting reliable digital data. Moreover, it is fundamental for automatic recognition within animal-based studies based on Artificial Intelligence.

## Introduction

Animal-based studies are crucial in approving pharmacological products and chemical substances for clinical applications [1, 2]. However, it is essential to prioritize the animals’ welfare in these studies [3]. Society widely recognizes the principles of animal welfare and the need to implement refinement, replacement, and reduction strategies (3Rs) when designing animal experiments [4]. Whether partial or complete, the replacement strategy involves utilizing well-established methods such as organ on a chip, organoids, or bioinformatical models [5]. While these methods can provide valuable insights into pharmacological interactions and the complexities of single organs, they cannot fully replicate the complexity of living systems [6]. Therefore, animal models are necessary to understand the underlying mechanisms comprehensively, particularly in areas like disease research, physiology, drug testing, and toxicity analysis [7]. To minimize the number of animals used in scientific studies, institutes and universities have established sharing programs for organs and tissues derived from animals. Additionally, training in Good Scientific Practice and properly handling laboratory animals enhances expertise, reduces the risk of experimental failures, and eliminates unnecessary repetition. Once the number of animals in a study has been reduced to a minimum, refinement strategies are implemented to minimize pain and distress and enhance welfare. Refinement includes environmental enrichment (e.g., running wheels, huts, toys) and adjusting the handling and housing conditions (e.g., temperature, noise) to reduce stress levels during experiments [8]. These refinement measures have proven to increase the value of acquired data sets. However, it is crucial to understand how human interaction and variability influence experimental outcomes and data sets’ reproducibility and impact clinical studies [9]. 1) Human handling and the presence of different individuals can induce distress levels in animals, even during routine procedures such as scoring and documenting severity levels. 2) The human subjectivity and interpretation of animal behavior and burden. To address this, digitalization using home-Cage-Monitoring (HCM) systems is necessary for documenting severity levels and minimizing human interaction.

Efforts have been made to digitalize animal studies by implementing home-Cage-Equal Monitoring Systems and additional monitoring tools [10–14]. However, there still needs to be standardized digital observation solutions for the actual home-Cage employed in biomedical and preclinical studies, which could further reduce human interaction as a refinement strategy [14]. In this study, we assessed the usability of a digital platform to monitor, record, and analyze laboratory animals in the actual Home-Cages to reduce human interaction as an effective refinement strategy for data acquisition and animal welfare. We developed a scalable monitor system and integrated it into the existing animal husbandry infrastructure at the Leibniz Institute of Virology (LIV) in Hamburg. The solution was implemented within an ongoing research project, and we conducted a complete proof-of-concept, including integration, service workflow implementation, and research and development, testing of hardware and software components. Furthermore, we examined the usage of the system during a common use case, post-operative monitoring. We analyzed the acquired data sets by using our software systems integrated review process, the reporting, and developed and implemented Machine Learning (ML) processes.

## Material and Methods

### Generation, Breeding, and Housing of mice

In this study, we employed parental female and male mice with transgenic NOD. Cg-Prkdcscid Il2rgtm1Wjl/SzJ background (Strain #:005557) from Jax Laboratory. Parental females and males were used to develop a breeding colony within the LIV animal husbandry. Mice individuals were observed within the study at a starting age of 10 weeks. To monitor the behavior level before and after surgery, we used mice individuals who underwent orthotopic transplantation of human liver tumor cells. The individual mice were marked and identifiable using ear punches or tail marks. Mice have been housed in individually ventilated cages (IVC) to exclude infection or transmission of infectious diseases. IVCs were changed as described in the standard operation protocol of animal husbandry.

### Surgery and Recovery phase

To achieve the goal of this study, we employed animals from an ongoing preclinical study (study approval number N56/2020). The goal was to monitor the mice before and after the transplantation of primary human tumor cells. Primary tumor cells were obtained from liver cancer resection (ethical study approval number PV3578; Amendment for this study was approved 18.06.2020; recruitment started from 01.03. 2021 and is still ongoing). Shortly, tumor cells were transplanted after isolation from fresh resections. The mice are anesthetized with isoflurane in all subsequent preparatory and surgical procedures. In the first step, the animal is injected subcutaneously with the painkiller metamizole (200 mg/kg). In the next step, a hair trimmer removes the abdominal hair from the animals. The shaved abdomen is then cleaned and disinfected with Betadine. This first cleaning takes place away from the operating table. After the onset of action of the painkiller metamizole (20-30min), the animal is now fixed on the operating table, and the abdomen is cleaned and disinfected two more times with Betadine. The operating table is tempered at 37°C to prevent the animals from cooling. A laparotomy is performed along the midline over a length of 3 cm. After imaging the left lateral lobe of the liver, intrahepatic injection of the tumor cells follows. Applying the cell solution preheated to body temperature (0,5×10E6 HCC tumor cells dissolved in sterile matrigel) is carried out in a standardized volume of 20µl utilizing a very thin injection cannula (30-gauge, 0.3 mm). Possible bleeding is quenched by absorbable hemostatic. After the closure of the muscles using a non-resorbable filament and skin using two clips, the mouse is separated from the isoflurane anesthesia and transferred to the conventional cage (Home-cage). Postoperatively, mice receive carprofen (5mg/kg subcutaneously) every 24 hours for 72 hours. The painkiller is stopped after 72 hours. Subsequently, the mice are examined daily to detect pathological changes. Pathological criteria include anemia (inspection of tail color), signs of local infection, splenomegaly (palpation), weight loss (weighing two times a week), as well as apathy and motor deficits (for example, sluggishness or pulling a limb).

### System architecture, hardware, and software

The iMouse solution is installed around existing home-cages via retrofit. Therefore, the hardware is named “DigFrame” because it encloses the home-cage. Our system consists of a standardized control unit per cage. This control unit is connected by 230V power and the internet. Within the control unit, we install one compute unit per camera. The camera (up to 5 cameras per Home-cage (front, left, right, back, top) is integrated into internally developed housings and integrated IR filter, equipped with a wide-angle lens of a view of 120mm. Cabling within compute unit and camera is done by a home-cage-specific pre-configured cable. The camera system is supported by a home-cage-specific, IR nightlight (920nm) to allow 24/7 remote observation [15]. To ensure that the field of view covered most of the home-cage, we set the camera at the four sides of the cage. Focus was on easy handling, simple installation, and serviceability. Cameras were mounted at the existing housing rack, located at the guide rails, which enables the same and consistent view in the IVC (**Figure 1A**). Each camera is connected to a dedicated computing unit. The computing unit connects to the computer network via a LAN cable connection. We explicitly decided to use a scalable open-source software named ZoneMinder as the foundation to observe the mice during their recovery phase and the experiment. ZoneMinder is a widely used and scalable monitoring platform system that can capture video from various camera devices and perform complex motion detection on the captured video in real time. We developed essential functions for the project and named the widely modified platform iMouseTV. The employed cameras were used for real-time monitoring, recording, and motion detection when setting up zones via iMouseTV platform. To record data, we implemented scalable computer hardware. This hardware was securely integrated into the existing IT infrastructure of the LIV. Users can access the iMouse solution by computer (full functionality) or the mobile app via tablet or mobile phone (view mode).

**Figure 1.**
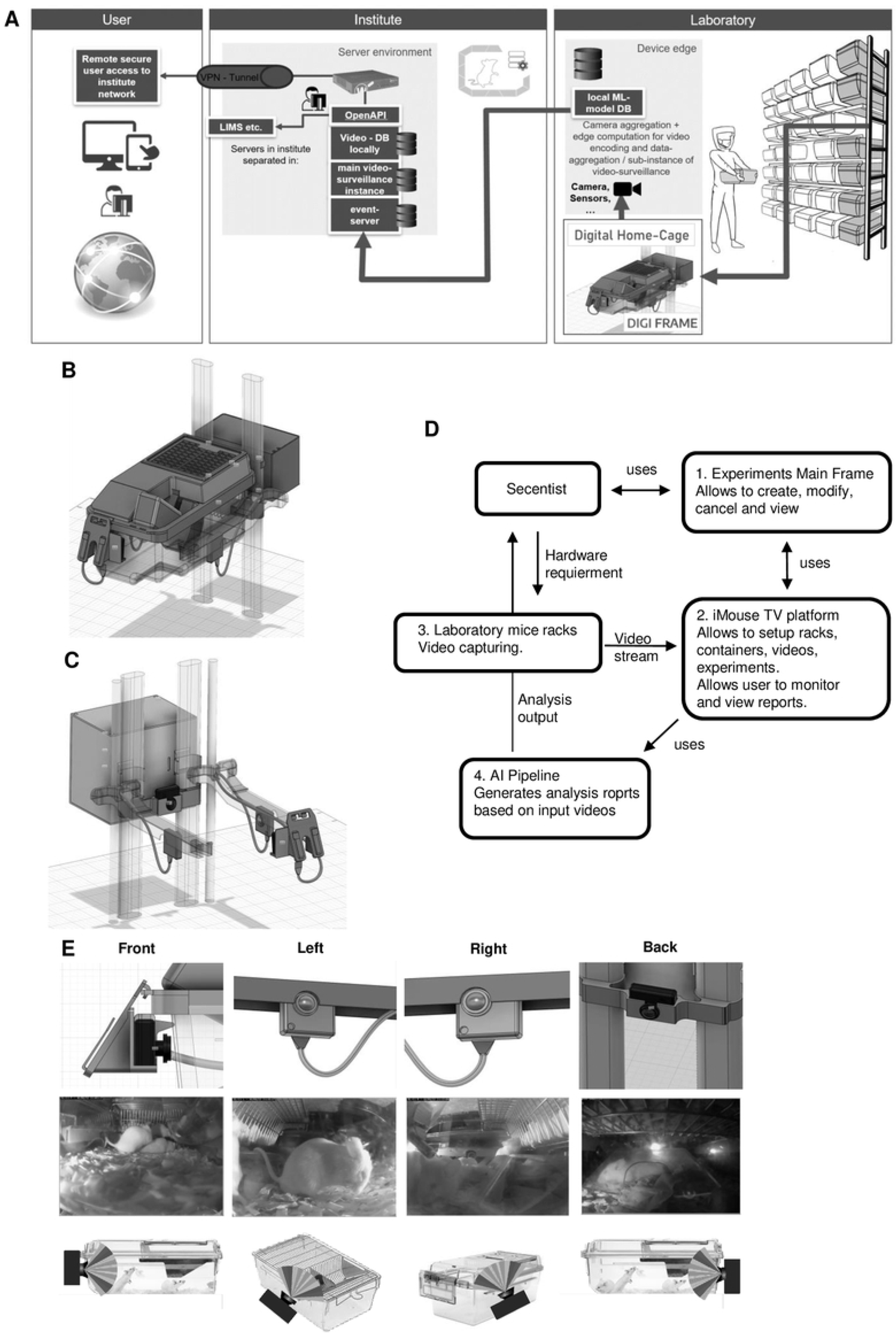
iMouse System Integration in the Existing animal husbandry Environment. (**A**) Illustration of the Security and Data Management Concept. The three main pillars are the laboratory, the institute network, and the user. The registered user can access the system remotely or within the institute network. The data collection and institute network are secured for remote access by implementing an OpenVPN. (**B-C**) Technical drawing of the Home-cage placed in the rack and the implementation of the observation units around the cage. Drawing showing the perspective with (**B**) or without (**C**) the Home-cage placed in the rack. Observation units are integrated at four positions, front, right, left, and rear (**B, C**). The observation units are connected to the Compute Unit behind the rack. The Compute Unit is connected to the institute’s electricity- and information network. (**D**) Simplified flow diagram of the iMouse system’s workflow, including the system’s Vision. The scientist or user will set-up the experimental frame (**1**), while the system will provide the data (**2, 3**) for the scientist and the AI pipeline (**4**). Here is the connection to the Laboratory management system possible (5). (**E**; upper part) showing the detailed technical drawing of the integrated cameras around the Home-cage. (**E**, middle part) Live view inside the Home-cage and the four perspectives of the cameras to monitor the different activity zones of the mice inside the Home-cage. (**E**; lower part) Representation of the four cameras and the focus on the Home-cage.

### Setting up zone-based recording

For the zone-based recording, we first created zones by naming them and adapting the zone shape to the spot of interest, e.g., the drinking outlet or the food area. Depending on the size of the zone and the predictable activity within the zone, we adapted the sensitivity of detecting, the change in the visible pixels or blobs in percent, and the method of alarm checking; both components have been used for starting the detection and recording of motions. E.g. for the detection of drinking activity, we used the type active; alarm pixel with a threshold of 20 (fast, high sensitivity), and for the climbing activity detection, we used the active type, with the alarm check method by measuring blobs, the threshold at 40 (best, medium sensitivity). To standardize the recordings with different zones, we implemented six shades of motion detection, three by blobs and three by pixel variation.

### Video recording and Storage

Recordings were performed for analyzing the zone-based activity level of the animals, for AI training, and for comparative analysis within the postoperative phase (7 days) of animals undergoing a surgery procedure. The recording of short video file recording was performed with a resolution of 640×480 pixels by 20 fps, and video files had a maximum length of 10s. For storage, the LIV internal server network and the LIV cloud storage for external access were used. Process-wise, all recorded video files were stored by date, camera name, and cage number to ensure traceability. For security purposes, all files were accessible only to authorized persons and encrypted in the storage space for additional protection.

### Video Processing by Learning Algorithms

The OpenCV framework within the Python programming language was employed to process the videos. The AI architecture relied on Pytorch, a Python framework for deep learning and processing unstructured data, and consists of 3 main models: YOLOv7 for object detection, ResNet50 for behavior classification, and DeepSORT for object tracking. The first two models demand data labeling as a precondition to train and implement them. This process is key as it allows the models to learn from the labeled data in a supervised learning process, as shown in **Supplementary Figure 3**. AI. The mentioned algorithm - based on the official repo [16] - is performed frame by frame on videos with the goal of detecting the desired objects. The second one enables tracking of the objects and differentiating between them, based on the objects’ previous and present positions and moves’ estimations. The last algorithm is used for behavior classification based on the cropped images of mice on each video frame. The goal is to detect mice in a video and classify the behavior of each of them. The output of such a process is in both textual and visual forms. The function returns a listing of information, such as which mouse and which activity, typically in a format called JSON (e.g., “labels”: [ “mouse1”, “mouse3”], “behaviors”: [“drinking”,” sleeping”], “coordinates”: [“0,3 100,200”, “400,100”,”500,600”]). The visual result is displayed as an image with the bounding boxes and labels drawn over the original image. The textual data is inserted into the database alongside the event record to be searched for and referenced later. The visual image is likewise stored alongside the event video for later display. Using 260 video events, 130 “real” drinking, and 130 “false” events, we reached accuracy levels of 89% in the drinking training phase, as shown in **Supplementary Figure 2**. After that, we used the longitudinal dataset for post-operation, as described in **Figure 4**, to tackle the ML algorithms.

## Results

### The digital units (DigiFrames) are implemented into the existing structure of the animal husbandry and the local IT-structure

Within the project, one of the main challenges was, integrating the iMouse solution into the existing structure of animal husbandry. Here, the concerns of integrating an open-source system for the facility’s users and operators had to be overcome first. For this purpose, we first developed a security concept that ensures the integrity of the research and the institute and protects sensitive and valuable data. This concept uses a VPN connection, as shown in **Figure 1A**. We started the implementation of the digital home-cages with just two DigiFrames on the right side of an ordinary rack system, which was used in an ongoing study on the LIV in Hamburg (**Figure 1A-C**). **Figure 1 (B and C)** shows that the observation and the control unit were integrated around the home-cages. They, therefore, didn’t change the handling processes of the facility employees or the user for the daily inspection or the changing of the cages. The control units are connected to the institute’s internal IT system via a LAN connection over a dedicated switch unit. A virtual machine (VM) hosts them on the institute server; the platform (iMouseTV) is accessed on the same VM. Videos recorded by the system are stored on the internal physical server, so the user can keep and review the files, including identifying metadata. Users can log into the iMouseTV platform via the institute network or, if not in the institute, via VPN, as illustrated in **Figure 1A**.

### The concept “DigiFrame” enables a complete view within the home-cage by covering several perspectives and all activities times

The goal of the underlying project was to generate a scalable monitoring system for home-cages in animal husbandry, providing users the full transparency of their ongoing experiments by displaying the day and night activity of mice in their natural habitat. Therefore, we established the DigiFrame, a retrofit adaptation on the home-cages, regardless of the manufacturing company. We designed the modular DigiFrame to be used for any home-cage. In this study, we show the concept of the DigiFrame representatively in one rack system, the Emerald line from Tecniplast. **Figure 1 E** shows we included four observation units (cameras) around the home-cage, with the resulting perspectives. As a result, the DigiFrame concept gives a complete view into the home-cage at any time since we also included a night vision system. The night vision also enables the simulation of day and night rhythms due to the use of a remote-control functionality via the iMouseTV platform. According to others, we utilized a specialized 920nm light-emitting diode (LED), chosen for its imperceptibility to rodents [15].

### The iMouseTV platform provides undisturbed, targeted, objective observation and reviewable and shareable raw data during and after animal experiments

In **Figure 1 D**, we show the simplified systems architecture. The user creates the input experimentally, while the recording will be managed automatically from the system. The user views the recordings and performs live observation for overall activity analysis or serenity assessment. Moreover, the recorded files were used for Machine Learning processes, and the algorithm produced reports for AI-based analysis as described previously. After logging in, the user can perform live view, recording, and motion detection, depending on the study and the needed dataset (**Figure 1 D**). As mentioned, we redesigned an existing, scalable, industrial-proven open-source software platform with scientifically necessary functions and created the iMouseTV platform with experimental focus. We implemented the possibility to allow and manage multiple user access simultaneously. After log-in, as shown in **Figure 2A**, the user attains directly to the specific dashboard. Here all experiments are listed, and the user can access the main functions of the system (**Figure 2 B-D**). The live view provides direct access into the observed and for the user assigned home-cages, as shown in **Figure 2B**. Here all perspectives are displayed. One of the main functionalities of the software is the experiment design, where users can set up the observation period of interest by setting up the timeline and the included observation units (cameras). Since the recording over a more extended period by several observation units leads to a large data set, we implemented a more specific recording option by creating zones of interest for motion detection (**Figure 2 C**). In Figure 2 C, two created zones are displayed in red, the Eating zone, and grey, the Nesting zone. Here the system provides different options for setting up the zones, as shown in Figure 2 C on the right side. Setting up the zone in the experimental setting is crucial for precise recording and preventing data waste. During a pre-study period of zone evaluation, we tested multiple conditions to determine the best options for being sensitive and accurate enough to detect all ongoing events but excluding events caused by light changes and reflections. Therefore, we implemented a pre-setup, listed in Table 1. We used the same settings for the same named zones during our study. **Supplementary Figure 1** shows the structure of the experiment section. The upper part is designed to give general information on the experiment, like name, duration, and owner. The user can see everything within the experiment, including observation units with their observation status. We also linked these thumbnails with the live view option. The recorded events are listed in the center of the experiment page, regardless of the recording method (zone-based or continuous). The system listed raw data for the events recorded during the set-up experiment on the right side of the experiment page. The events are plotted time-dependent by the hour per day and duration in seconds. In **Supplementary Figure 2**, a representative event is shown. The systems provide all event metadata information, leading to distinct traceability and transparency of all events.

**Figure 2.**
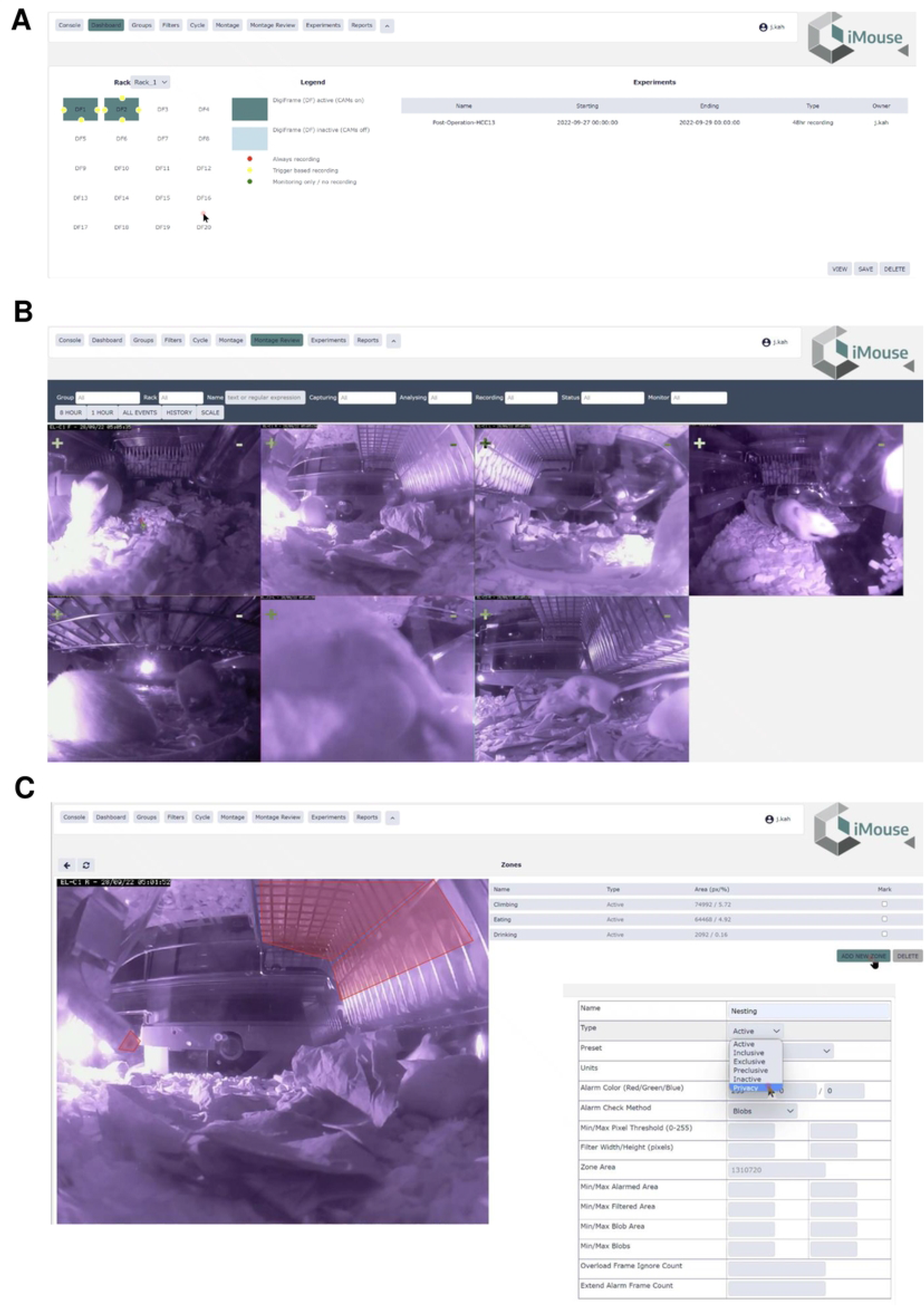
Description of the main iMouseTV platform functionalities. iMouseTV platform screenshot, showing the main functions necessary to set up, view and analyze experiments. (**A**) Displaying the Dashboard of a registered user. The available observation units (physical camaras) for the user are displayed on the upper part. The user can choose the husbandry rack and the DigiFrame used for the ongoing experiment. The observation units are shown as dots around the cage; here, the amount, the position, and the recording/ monitoring status are mentioned (Legend). The lower part of the dashboard shows the user-specific experiments, which can be viewed and edited by the user. (**B**) Showing the live view page of the system. The user can choose a particular DigiFrame for observation. (**C**). Displays the Motion Detection option of the system. The user can create zones of interest from every observation unit, specifically for every experiment. The user can name and fit the zones to the area of interest.

### Motion detection-based recording enables multiple activity analyses during post-operation monitoring

To evaluate the relevance of accessing a home-cage monitoring system like the iMouse solution during preclinical basic research studies, we employed NSG mice, which are used for orthotopic transplantation of liver cancer cells, to establish a therapeutic testing model of liver cancer drugs. For the general analysis, we observed 3 NSG mice in their home-cage. The three female mice were hosted in the same cage for three days after maintenance and before operation. After the operation, the mice were returned to their familiar home-cage for the wake-up phase. After the wake-up stage (20min), the mice were placed back into their racks, into the DigiFrame slot for observation. The user started the pre-set-up experiment and ran the experiment for seven days under the same litter and enrichment conditions (no houses, no toys, no tunnel, but cellulose). The reason for the low enrichment is the usage of metal clips for wound closure. The system recorded the motion in the climbing, eating, drinking, and nesting zones. As shown in **Figure 3 A**, the system recorded activity levels higher at night and lower during the day. The system summarizes the activity duration and displays those per hour and day. Therefore, **Figure 3 A** plots a higher activity level as a longer duration. For reviewing the recording and explaining the recording modalities given by the system, we selected a single event, 731553, which was detected on the 19^th^ of Dec 2022 at 08:13:00 by the observation unit EL-C1 R (pointed out by the black arrow in **Figure 3 A**). The event includes a high number of alarm frames, which are visible in the lower timeline (red columns) and refer to multiple activity recordings. As shown in the single frame **Figure 3 B**, all three mice are active in different zones. Therefore, the reason for motion detection is climbing, drinking, and nesting. To describe the complexity of the motion-based recording, we analyzed two single frames (ID 116 and ID 120), shown in **Figures 3C and D**. Both frames reached the threshold for setting an alert and therefore recording. The single frame 116 scores 40 as an additional effect from the drinking zone pixel changes (16%) and the climbing zone blob changes (30%). The single frame 120 scores 35 due to the climbing zone blob change. This is in line with the visible movement of the mouse in the climbing zone in frame 120 (marked by black arrows), compared to its position in frame 116, which results in the recorded blob changes of the analyzed single frame. In comparison, the mouse in the drinking zone is causing a lower change in the pixels (9%), which is below the detection limit (10%). Therefore, when setting up multiple zones for activity recording, distinguishing single activities for single mice is not viable. Nevertheless, motion drinking was part of both alert frames. Therefore, the event can be used as a “real” event for displaying the occurrence and the frequency of the drinking event. It can provide activity measurements without disturbing the mice during their recovery and post-operation phase. The recorded dataset shows a high activity level shortly after the operation and points out that the operated animal recovered much earlier than expected. Therefore, the more intensive observation period could be reduced from 72h to 24h for this model to lower human interaction and stress induction.

**Figure 3.**
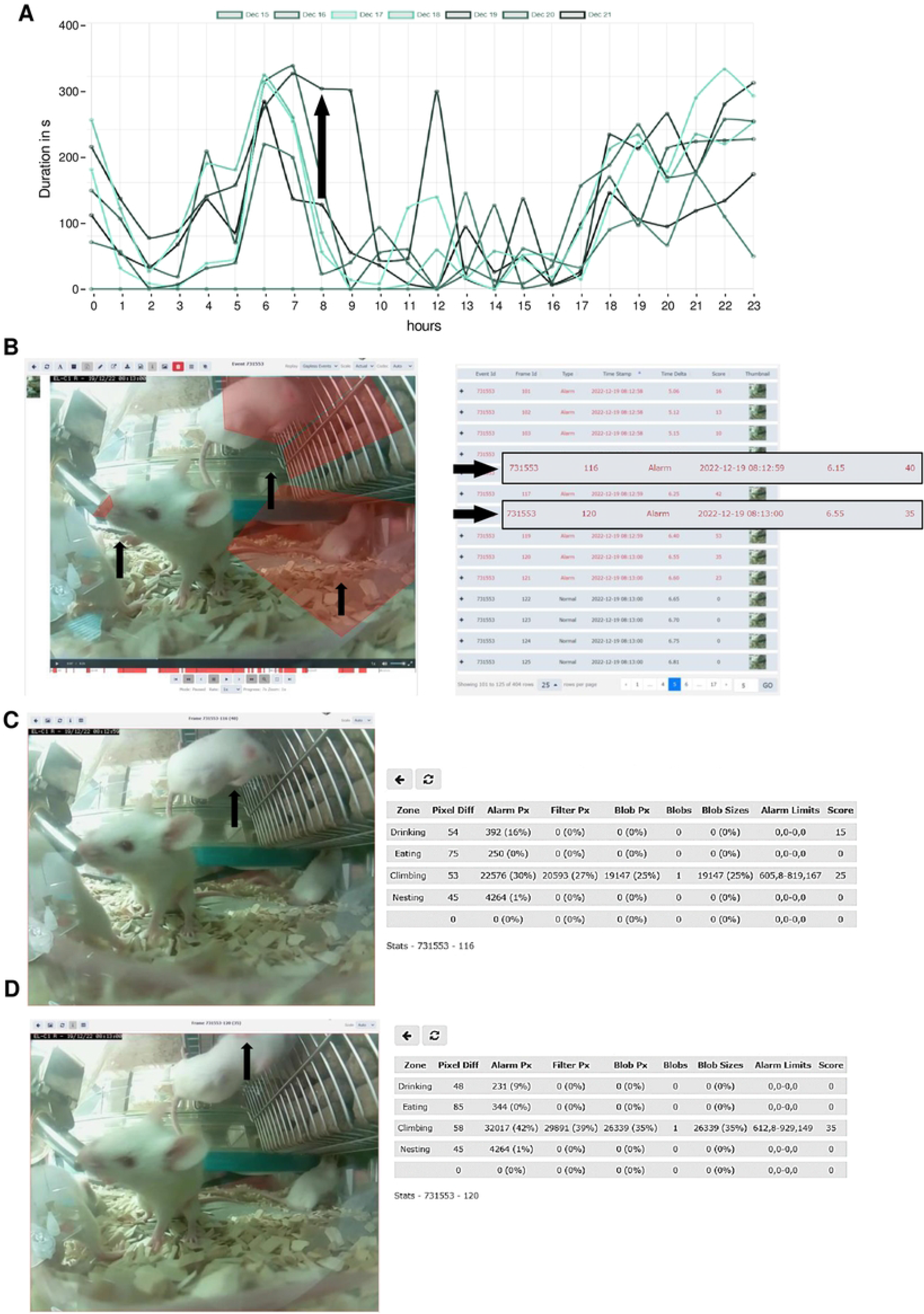
Motion detection and activity recording during postoperative monitoring. (**A**) Activity raw data analysis provided from the iMouseTV platform after seven days of postoperative monitoring of one DigiFrame using motion detection. X-axis plots the hours per day; on the y-axis, the duration of the motions during the day is visualized. The days are displayed from the system in different colors and mentioned in the upper part of the graph. (**B**) Single frame extraction of one motion detection plotted in **A**, visualized by an arrow pointing to the frame’s metadata. On the right side, the single frames are listed in a table. The frame table gives information on the event ID, the Frame ID, the type, the time stamp, the time delta, and the score for every frame. The fundament for the motion analysis, the type, the time delta, and the score are crucial. On the right side, two single frames (Frame ID 116 (upper frame) and 120) are pointed out by black arrows. The system types both frames as alert frames. During the experiment, motion detection was performed for four zones (**C and D;** eating, drinking, nesting, and climbing). In (**C**), frame ID 116, and in (**D**), frame ID 120 are displayed, including the system’s motion detection analysis raw data table (on the right). The raw data table shows the pixel differences in all four experiment zones. **C** shows that the system mentioned alarm pixels in the climbing (30%) and the drinking (16%) zone, giving an overall score of 40, as also mentioned in **B**. In D, the alert table displays the alarm pixels in the climbing (42%) and the drinking (9%) zone, leading to an overall score of 35.

### Machine Learning algorithms enable the system to identify individuals and their behavior without human bias

To understand the potential, the advantage, and the limitation of implementing Machine learning algorithms within digital observation, we reanalyzed the longitudinal post-operation dataset **(Figure 3)** using AI models. We used ML algorithms to analyze the video files acquired by motion detection. The videos were recorded in both, day, and night modes. The first step was to randomly extract the drinking events from the overall activity levels of mice after the operation located in the cage for seven days. After the extraction, we divided the dataset into a training set and a validation set – during the later stages, data extracted from the specific video will not be reshuffled but will belong to the same set as the video; that would prevent data leakage between the training and the validation sets. Next, data preprocessing and labeling steps were taken as a precondition to implement the ML models, as summarized in **Supplementary Figure 3**. **Figure 4** illustrates the outcomes of the AI modeling, encompassing mice detection, behavior classification, and visual object tracking. **Figure 4** is therefore divided into Metrics describing the statistical results of the modeling part; the output bar chart generated after analyzing a short video file and the visual confirmation of the achieved results. The first part presents metrics that describe the statistical results of the modeling process. **Figure 4 A** represents a confusion matrix displaying the classification outcomes on the validation set. This matrix provides detailed information about the classification tasks, going beyond simple accuracy measurements. For classification, we utilized 1167 cropped frames of mice, with a roughly equal distribution between daily and night modes. The dataset comprised 21% positive class images (’drinking’) and 79% negative class images (’not drinking’). The training set constituted approximately 83% of the dataset, while the validation set accounted for 17%. We employed the oversampling method to balance the classes, which yielded favorable results. Ultimately, we achieved an accuracy of 94.36%, with a sensitivity metric of around 88.1% and a specificity metric of approximately 96.08%. It can be anticipated that as the dataset’s classes become more balanced, the disparity between sensitivity and specificity metrics will diminish. **Figures 4B and 4C** display the general outcomes of the object detection modeling results. **Figure 4B** presents the overall results of the process of training the object detection model evaluated on the validation set. The chart displays the mean average precision metric calculated at IoU (Intersection over Union) threshold amounting to 0.5 (‘mAP0.5’), precision metric, and recall metric concerning the iterations during the training process. **Figure 4C** presents the precision-recall curve displaying the tradeoff between precision and recall for different threshold levels measured after the last iteration of the process of training the object detection model is completed. The precision-recall curve was built based on the validation set. The area under the precision-recall curve yields the same result as the ‘mAP 0.5’ measured at the end of the training process. We relied on 708 video frames for the object detection component, with a comparable number of images in both day and night modes. The training set encompassed 82% of the dataset, while the validation set constituted 18%. Our conclusions were based on the previously mentioned ‘mAP 0.5’ metric. We achieved an approximate mean average precision score of 0.819, enabling the successful detection of mice in both daily and night modes, forming the foundation for our subsequent analyses. **Figure 4D** shows the output data in the form of a bar chart. This chart displays the total number of mice detections categorized by their behavior and unique ID numbers. It was generated after analyzing a short video comprising 210 frames. It displays the individual behaviors of the three mice hosted in the cage and can differentiate that mice ID1 and two were not drinking while the mouse with ID3 was drinking. **Figure 4E** showcases visual confirmation of mice detection. The AI algorithm generates predictions once the confidence level reaches a certain threshold (in this case, 49%). Each detection is visually demarcated by a bounding box, with the confidence level displayed in the top-left corner of the box. **Figure 4F** illustrates the visual confirmation of object detection and tracking. Each tracked object is represented by a red dot positioned at the center of its respective bounding box and a unique ID number adjacent to the center point. The tracking of mice relies on their previous and current positions. Finally, **Figure 4G** combines object detection, tracking, and behavior classification components. The unique ID number of each mouse is displayed in the top-left corner of the frame, accompanied by the predicted behavior for that particular mouse.

**Figure 4.**
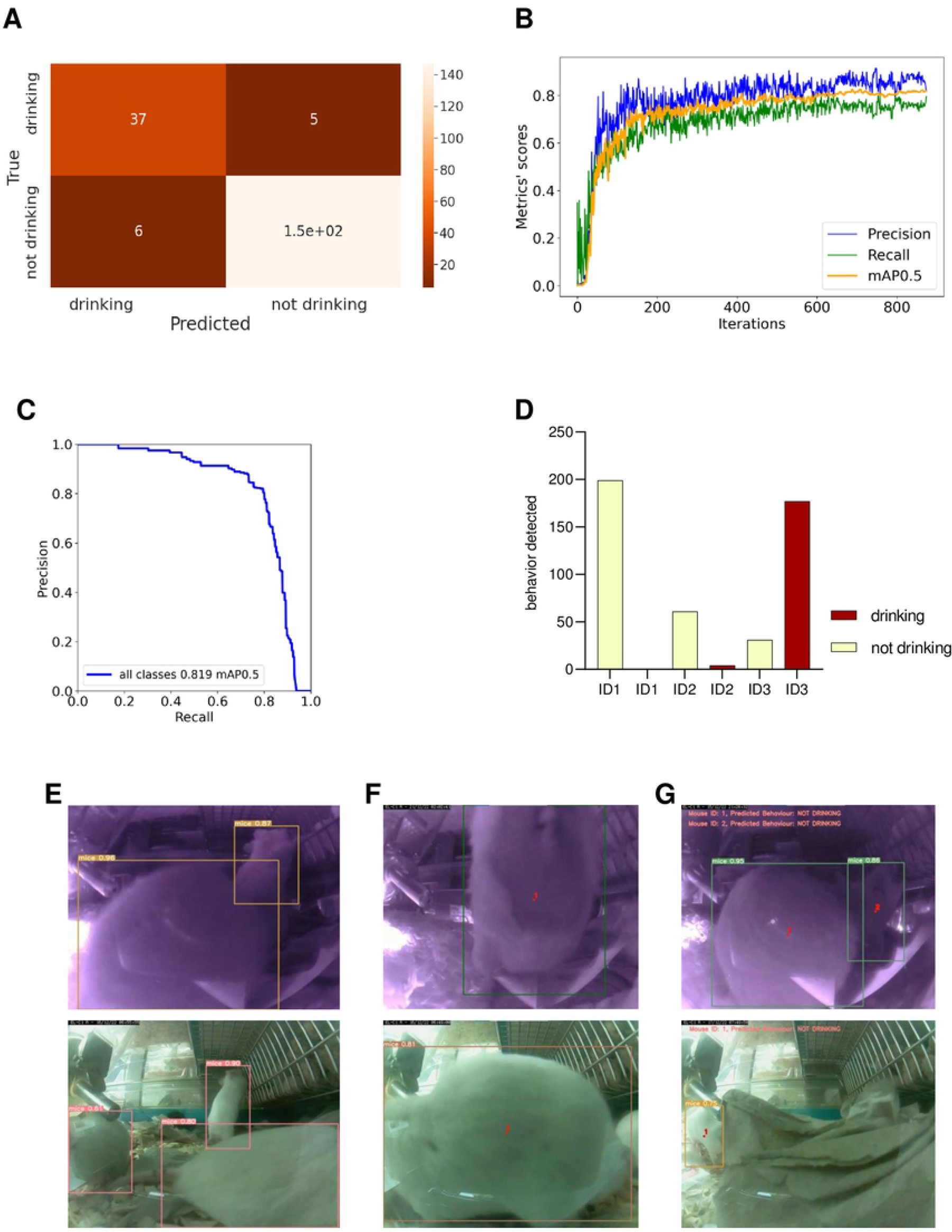
Machine Learning training enables analyzing drinking behavior. Visualizations show metrics displaying the results of the modeling part (**A, B, C**), an example of the output data plotted in the bar chart (**D**), and examples of the achieved results in the form of information placed on the extracted video frames (**E, F, G**). (**A**) Confusion matrix displaying the results of the behavior classification on the validation set. Upper-left side shows true positives, the upper-right shows false negatives, the lower-left shows false positives, and the lower-right shows true-negatives. (**B**) The plot shows the precision, recall, and mAP0.5 scores during the training process in relation to the iterations. Y-axis shows the metrics scores, and X-axis displays the iterations. (**C**) Precision-recall curve determining the tradeoff between the precision and recall for different threshold levels. Y-axis shows the precision score, and X-axis shows the recall score. In (**D**), results of individual mouse drinking/not drinking behavior were plotted, displaying three mice recorded within 10s at 10 pm on Dec 15. The bars represent the number of events detected in relation to the predicted behaviors on a random video. Y-axis shows the number of detections, and X-axis displays mouse ID in one cage simultaneously. In (**E-G**), representative visual confirmations of object detection, behavior classification, and object tracking parts are shown. Extracted frames in night mode are displayed on the upper and lower sides in day mode. (**E**) shows the visual confirmation of the object detection with bounding boxes around each detected object. The object’s class names and detection probabilities are displayed on the top-left side of each bounding box. (**F**) shows the combination of the object detection and object tracking parts. A red dot informing about the center point of each detection is placed on the center part of each detected object with the unique ID number just next to it. (**G**) displays the combined results of object detection, object tracking, and behavior classification. The predicted behavior of each detected object is displayed on the upper-left side of each frame in the form of short information containing the predicted behavior and the object’s ID number.

### Implementation of the Artificial intelligence system refines motion detection-based drinking dataset for the post-operation experiment

To acquire data for a specific activity from a series of zone detection experiments, we utilized machine learning (ML) algorithms. In **Figure 5**, we initially filtered the recorded events from the same experiment based on the “Motion: Drinking” category, and the resulting data is presented in **Figure 5A**. The peak observed on December 19th (Monday) can be attributed to the weekly cage-changing activity. Since this particular analysis aimed to evaluate the accuracy of our method, we opted for a manual evaluation of drinking events. To streamline the review process, we reduced the number of events to be assessed from 7 days to 2 days. **Figure 5D** displays the results after human experts reviewed the events. We then implemented the ML algorithms based on the events extracted from the iMouseTV platform and evaluated the results. **Figure 5B** displays the performance of the ML algorithms over the period of 7 days. Then we decided to reduce the analyzed period to compare it with the period reviewed by the human experts. **Figure 5C** displays the performance of the ML methods on the period of selected two days based on the overall data extracted from **Figure 5B**. Then it became possible to compare the performance of the ML algorithms (**5C**) with the results obtained after the human experts (5D) reviewed the events. During seven days, the AI system recognized mice drinking behavior in 12.46% of the video frames that had been previously filtered using the system’s “motion: Drinking” behavior filter. Additionally, the system detected mice drinking during 6.94% of the instances they were detected overall. The AI system confirms the peak of mice drinking activity on December 19th. Generally, mice were observed to drink most frequently during evening hours, with significantly reduced drinking activity during the day. Utilizing AI algorithms enables us to enhance the accuracy and efficiency of the analysis.

**Figure 5.**
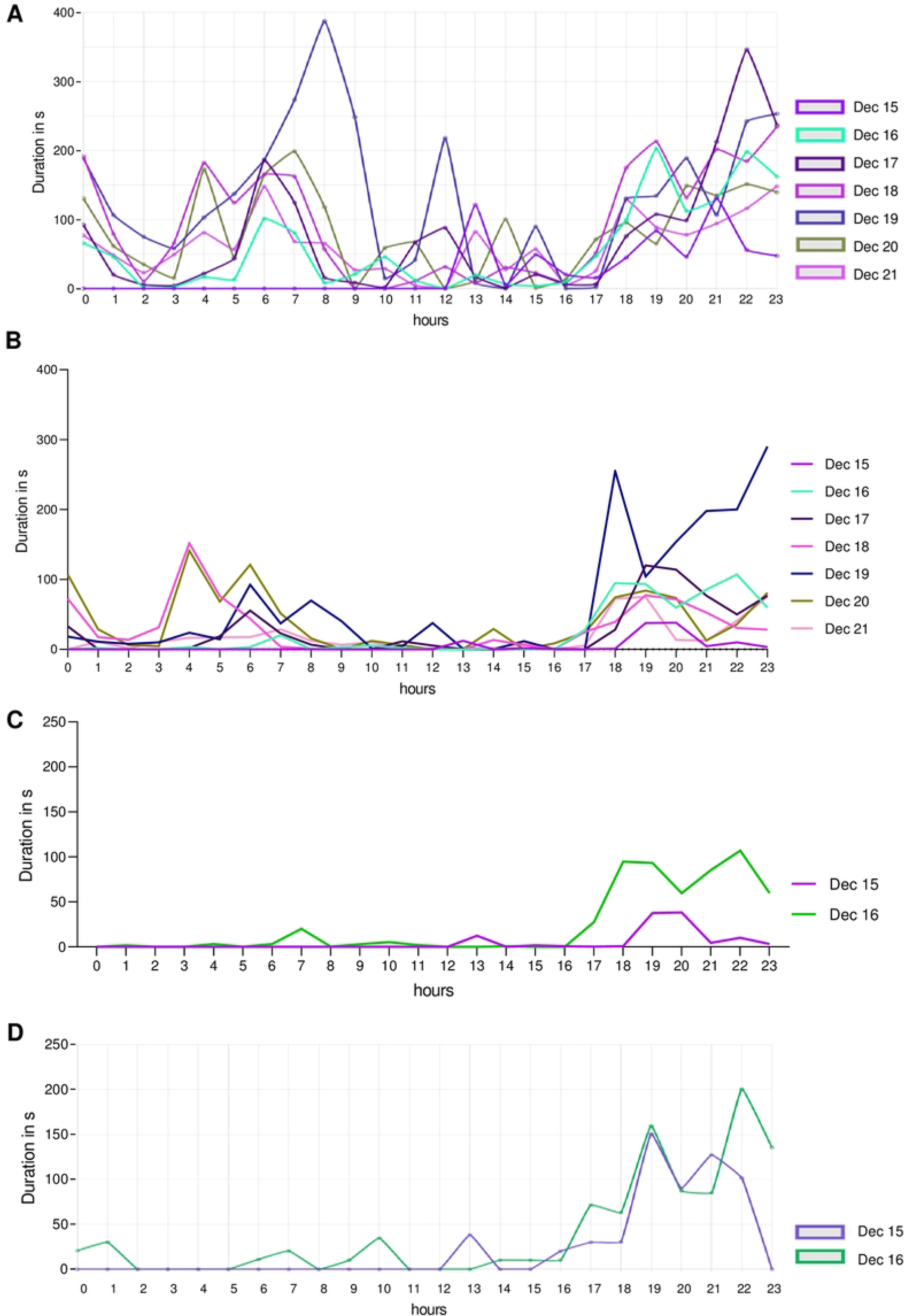
Comparative detection for drinking events during post-operation using a systems-based report system and trained ML algorithms. Time-depended duration of detected and recorded drinking events, spanning from the recovery phase till one week after transplantation of three mice in their Home-cage. Results are plotted in duration in seconds per recorded event against the hour. The human-based “by-hand observation” has proceeded as described in the animal experiment approval (N056/2020). (**A**) Detection for the drinking zone, extracted from the iMouseTV platform. (**B**) ML algorithms pre-analyzed data set. (**C**) ML pre-analyzed data set of 36h after transplantation, extracted from (**B**). (**D)** Represents the plotted data of the first 36h after transplantation reviewed in the sense of “false” and “real” event by the user.

## Discussion

Society and the scientific world are facing ever-increasing problems related to animal experimentation. These include the reproducibility crisis and the general use of laboratory animals in preclinical and clinical research [14, 17, 18]. The fact that animal testing cannot be replaced entirely and is a crucial part of translational research becomes clear with introducing of new drugs and chemical substances, which the legislator determines [2, 19]. Also, significant efforts have already been made to replace animal experiments, which have already achieved great success and significantly contributed to solving one of the problems. Nevertheless, due to the complex systemically effective drugs, their effect can only be mapped in the living organism. Physiological relationships and immunological issues can also only be investigated in depth in systems that provide a multi-organ basis. This is where the methods of animal reduction reach their limits, and the amelioration methods for animal experiments take effect. We know that refinement of animal experiments goes beyond simply adding more nesting material. At this point, improving data acquisition and utilization takes a leading role [20, 21]. In preclinical research studies and in the pharmacological industry, monitoring and data collection is mainly performed by human beings during the daytime, with their subjectivity and the fact that most animals in studies are mice, which are night-active and fleeing animals by nature. This kind of methodology is fundamentally wrong. Here we need to improve preclinical studies, and we still face a technology limitation for monitoring techniques available for animals in experimental settings. Actual methods are limited in precision, scope, or invasiveness [22]. These issues present challenges for researchers in obtaining high-quality data sets. We already understand that new methods and strategies must be worked out to improve data acquisition [21, 23, 24]. To refine research experiments, collecting a higher amount of data points and additional information having an impact on the interpretation of the results is crucial. At this point, we receive large data sets, therefore seeking the next challenge, handling and interpreting those big data sets [25]. Implementing Machine Learning and Deep learning will give rise to the collected data set. The advantages of ML were used in the fields of translational and medical research for screening and analyzing patient-derived data sets [26]. In genomics and neuroscience research, as well as in the prediction of zoonotic animal movement [27], ML processes are used to accomplish research hypotheses and, as recently shown, to predict and classify animal motions without human interaction [28]. We believe that reducing human bias in preclinical studies can improve the reliability and quality of results and increase animal welfare.

Therefore, we investigated the use of a camera-based monitoring system installed around IVC cages to evaluate the feasibility of visual non-invasive monitoring in combination with recordings of this monitoring. Issues of recording quality, manageability of data, traceability of data, and integrability of the system within the animal house were first evaluated. Subsequently, we focused on the data analysis, here, we investigated the reduction of the amount of data and the use of the processed data for the training and finally for analysis using a well-trained algorithm cascade. As a result, we could show in this study that the iMouse solution can be integrated into the research environment. We showed that using the system by the scientific staff within the institute and remotely is possible. The data handling and analytics process, including metadata and time stamps, were generated for traceability and assignability of raw data, assignment by name, date, and time. The digital observation platform was adapted within the study to allow the user to set up single experiments and view and evaluate it separately within the system in an initial overview evaluation. Various monitoring methods are possible within the system, from manual observation of the single cage via live view to recording a specific period.

In most cases, only certain behaviors of the animals are informative for observation, such as the frequency and duration of a drinking event. For this, we found a feasible solution through the system by marking a relevant area (zone) to be used as a reference for initiating a recording. A video was recorded if a change in image composition was detected in this area. However, if within one observation, several areas, e.g., also feeding or climbing, are of interest, the system could ensure a multi-zone recording, but due to the proximity of the zones, here only the evaluation as total activity was possible.

To ensure further separation of individual behaviors from a multi-zone recording, we used the possibility to evaluate these total activity events by using self-trained algorithms. We could show here that this kind of processing is valuable and feasible. Through this, we could achieve a further significant increase in the precision and quality of the data.

In sum, we established a standardized camera approach by using the common home-cage as the fundament and therefore tackle the missing link between animal-based preclinical studies and the achievement of unbiased data sets. Moreover, we implemented a data reduction and refinement strategy for the acquired data sets using zone-based detection. Subsequently, we successfully integrated ML processes for individual mouse recognition and behavior detection using iMouseTV platform recorded data sets.

For the first time, we showed a complete digitalization approach for the preclinical research field, achieving unbiased data within the common mouse home-cage. The need to improve data collection is becoming increasingly important, especially when it comes to the approval of new drugs. Here, the failure rates of individual substances are very high, especially regarding efficacy and safety [29]. Using an unbiased preclinical measurement of effectiveness and safety can reduce these failure rates, as these lay the foundation for initiating a clinical phase II and, ultimately, phase III trial.

### Outlook

Our upcoming efforts will concentrate on three main parts. Firstly, the iMouseTV platform improvements regarding user management and functionality. The overall system optimization includes scalability, data usage, and handling. And thirdly, the increase of AI and Machine Learning accuracy, efficiency, and amount of recognized behaviors, meaning higher prediction level using fewer data sets. Our overarching objective is to train ML algorithms using a community-based model that links interconnected laboratories. This approach aims to elevate data generation and utilization quality, accuracy, and reliability.n

### Study approval

Primary human cancer cells were isolated from resections patients suffering from HCC using protocols approved by the Ethical Committee of the city and state of Hamburg (PV3578) and accorded to the principles of the Declaration of Helsinki. Patients consent to the study in written words. Animals were housed under specific pathogen-free conditions according to institutional guidelines under authorized protocols. All animal experiments were conducted by the European Communities Council Directive (86/EEC) and were approved by the City of Hamburg, Germany (N056/2020).

## Author contributions

JK, ML initiated and supervised the research study; IC designed and remodulated the software, DS, ML, and ML designed the hardware. DS and ML integrated the hardware. JK, ML, MF, UM designed and conducted experiments, and JK, acquired data. IC, MF, and JK analyzed data; JK, ML, NS, and OS wrote the manuscript. All authors discussed the data and corrected the manuscript.

## Acknowledgments

We thank Tobias Gosau for this excellent work with the mouse colony. We thank Norbert Zangenberg and Heiko Juritzka for their IT support during the project. We thank the LIV administration for the fruitful cooperation.

## Financial supports

DFG founded the animal experiments (KA 5390/2-1). Hardware and Software were financed by IIoT-Projects GmbH, private investors, and Thaumatec Tech Group.

---

**Supplementary Figure 1**

***Presentation of the experimental page in the systems platform.***

Shows the Experiment page of the system. The system acquires events and collects them into an experiment folder. On the right side, the first day of an experiment is displayed, showing the zone-specific events in time on the x-axis plotted against the duration of the events in sum on the y-axis. All observation units being part of the experiment are listed on the left side. The listing here is also categorized into the DigiFrames.

**Supplementary Figure 2**

***Descriptive presentation of a recorded event.***

Displays a single frame of a recorded event (video file) in the Home-cage showing experiment-specific zones for, e.g., drinking, climbing, nesting, and eating, marked by white arrows. Also shown is the meta-data implementation (Name of the Observation unit, date, and time of the recording) at the top, and on the left are the properties listed bellowing to the event. The meta-data shown in the video file does allow unambiguous assignments. In the lower part, the timeline for the video is displayed. Within the timeline, the activity peaks are shown in red columns. The properties of the video, e.g., the storage path, the time, name, and duration, are uniquely assigned to the video and are shown in higher magnification in the lower left. The recorded videos can be played at different speeds to watch specific alerts. The system records the events in single frames. Therefore, the user can watch and analyze the event from frame to frame.

**Supplementary Figure 3**

***Training of Machine learning algorithms.***

In (**A**), data labeling and annotation were displayed with CVAT and Roboflow. Most of the labeling process is based on the Roboflow framework that facilitates open-source functionalities. Roboflow itself is a developer framework focusing on Computer Vision. It enables quick and practical steps covering data collection, preprocessing, and model training techniques. (**B-D**) Shows the export data generated in “YOLOv7Pytorch” format— the output in the analytical form represented by the three charts. (**B**) The graph represents the number of frames each mouse detected on the video. (**C**) represents the general drinking statistics. It represents the sum of frames that all the mice were spotted drinking or not drinking. It amounts to the combination of frames representing several mice. (**D**) presents detailed drinking statistics. It represents the number of frames that each mouse was spotted drinking by the unique ID number of each of them. (**E**) represents output data for the single frame shown in (**A**), including the mouse ID and the probability of the behavior.

